# StrucMind links protein contacts and ligand environments to atoms in PyMOL

**DOI:** 10.64898/2026.11.26.769536

**Authors:** Manal A. Nael, Laxman M. Alakonda, Khaled M. Elokely

## Abstract

StrucMind provides two PyMOL plugins for comparing protein contact maps and examining the interactions around a bound ligand. A contact seen in only one structure can reflect a difference in packing or a gap in the coordinates needed for comparison. Contact Maps aligns corresponding residues, identifies retained, gained and lost contacts, and records why a comparison remains unresolved. Structure Analysis measures direct ligand–protein contacts, traces paths through individual water molecules, and characterizes aromatic packing and pocket geometry. Users can select a contact or a water-mediated path to inspect the atoms in PyMOL. Exported atom identities, distances and coordinate coverage let a colleague retrace the assignment.

---

StrucMind links each contact on a residue map to the atoms used to measure it in PyMOL [1]. A missing segment in a deposited structure can leave an apparent contact change unresolved [2]. To interpret a gain or loss, a researcher needs to know both where the atoms are close enough to form a contact and whether the other structure contains the atoms needed to establish its absence.

Both plugins run locally. In Contact Maps, users select the chains or assembly copies, designate a reference structure and choose a contact type. For heavy-atom packing, they also set the distance cutoff. Selecting a residue pair displays the measured atoms in each available structure. The Partner changes view lists the partners gained or lost at each position. Structure Analysis starts with a ligand copy and the selected receptor chains. Users can inspect a direct contact or follow a path from a ligand atom through a water oxygen to a protein atom. The exported path identifies all three atoms and gives the distance for each leg.

We first asked which contact differences between two human dihydrofolate reductase structures (DHFR, EC 1.5.1.3) could be evaluated in both. The folate-bound structure 1DHF and the methotrexate/NADPH-bound structure 1U72 share chain A residue numbering, including positions with missing coordinates [3,4].

We generated one chain A packing map per DHFR structure with each of two tools, Contact Map Explorer 0.7.0 [5] and native PyMOL 3.1.0. Both enumerated residue pairs with occupied heavy atoms within 4.50 Å, including sequence neighbors. For each tool, we matched the two maps by deposited residue identity (Figure 1). Both workflows yielded 811 pairs in 1DHF, 843 in 1U72 and 777 in both. This left 34 pairs observed only in 1DHF and 66 only in 1U72. Data S3 supplies both scripts and all 877 pairs.

**Figure 1.**
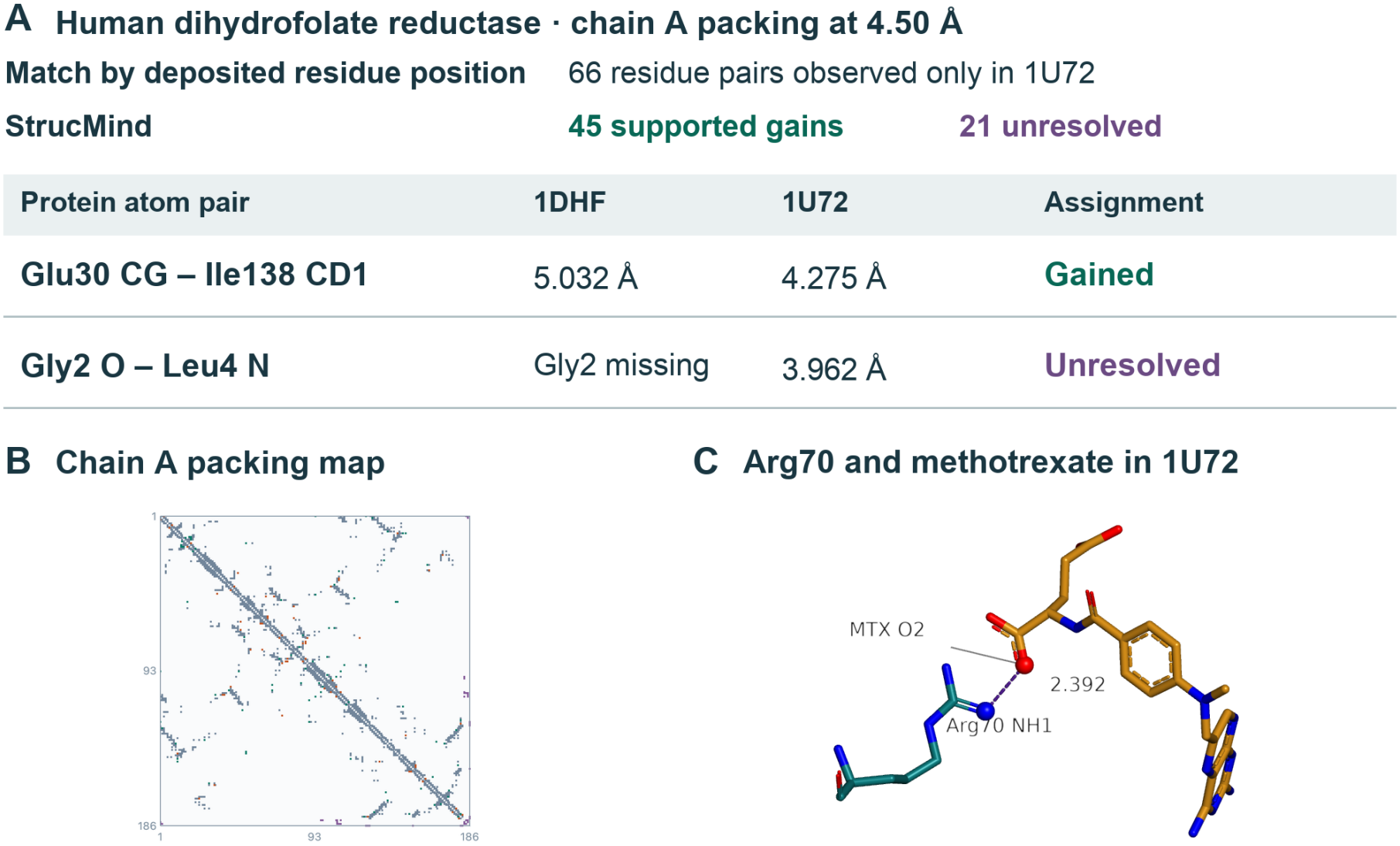
Chain A of folate-bound 1DHF and methotrexate-bound 1U72. (A) Each program generated a separate packing map per structure. Matching pairs by deposited residue identity yielded 66 observed only in 1U72. Glu30– Ile138 crosses the 4.50 Å cutoff. Missing Gly2 leaves the reference status of Gly2–Leu4 unresolved. (B) The contact map shows 877 pairs. Teal denotes gained, orange lost, slate retained and purple unresolved. (C) The PyMOL view shows Arg70 NH1 and methotrexate O2 at 2.392 Å.

Of the 66 pairs observed only in 1U72, 45 qualified as gains. For example, the closest heavy atoms of Glu30 and Ile138 were CG and CD1 in both structures. Their distance fell from 5.032 Å in 1DHF to 4.275 Å in 1U72, crossing the packing cutoff. The other 21 pairs involved at least one of four residues missing from 1DHF (Val1, Gly2, Ser3 and Asp186). Gly2 O is 3.962 Å from Leu4 N in 1U72. StrucMind preserves that measurement and records why the reference distance is unknown. All 34 pairs observed only in 1DHF qualified as losses.

An observed atom pair can establish packing in a partially resolved standard residue. A supported absence requires unambiguous mapping and sufficient coverage of the expected heavy atoms. The export distinguishes missing positions, incomplete residues and alternate conformers (Supporting Information, Table S1 and Data S1–S3). Selecting methotrexate A188 in 1U72 illustrates the corresponding ligand workflow. Among 203 ligand–protein atom pairs across 19 residues, selecting the closest one displays Arg70 NH1 and methotrexate O2 at their measured separation of 2.392 Å (Figure 1C).

CMView displays shared and unique contacts on structures [6], RING-PyMOL compares residue interaction networks [7], and difference maps can exclude unobserved residues [8]. A StrucMind export retains the observed atoms even when the opposite structure lacks the coordinates needed to evaluate the pair.

Across six deposits, a separate spatial enumeration reproduced all 49,443 protein packing pairs. We remeasured 78,365 atom-distance records and both legs of 3,234 water-bridge records (Data S1). The Supporting Information documents the verification scripts, input accessions and expected measurements.

The ligand analysis asks which protein atoms a ligand contacts directly or through water. Arpeggio and PLIP identify atom-level interactions [9,10], LigPlot+ compares interactions across related complexes [11], and ProLIF retains the atom indices behind interaction fingerprints [12]. PLIP also reports the water and both distances in a water bridge [10].

A water near a ligand can also make polar contacts with the protein [13]. LeView depicts water-mediated paths in ligand–protein interaction diagrams [14]. The carbonic anhydrase II structure 3HS4 (EC 4.2.1.1) contains three acetazolamide copies [15]. For active-site AZM A701 (segment D, model 1), 94 atom pairs link the ligand to 13 chain A residues within 4.5 Å. Ligand N1 lies 2.825 Å from Thr199 OG1. Which of the three deposited waters within 4.0 Å of this ligand also contacts the protein?

Only water 1191 connects the ligand to the protein under the geometric rule (Figure 2). Four ligand atoms reach two protein atoms through that water, yielding eight paths. Users can keep all eight or select the two that meet the polar rule (N or O at both endpoints, both distances ≤3.5 Å and water occupancy ≥0.70). Both polar paths start at acetazolamide N2, 2.837 Å from water 1191 O. The same oxygen lies 2.754 Å from Pro201 backbone O and 2.906 Å from Thr200 OG1. The export therefore identifies one bridging water with two candidate polar contacts on the protein side. Independent measurement reproduced all 94 direct ligand–protein distances within 0.0001 Å.

**Figure 2.**
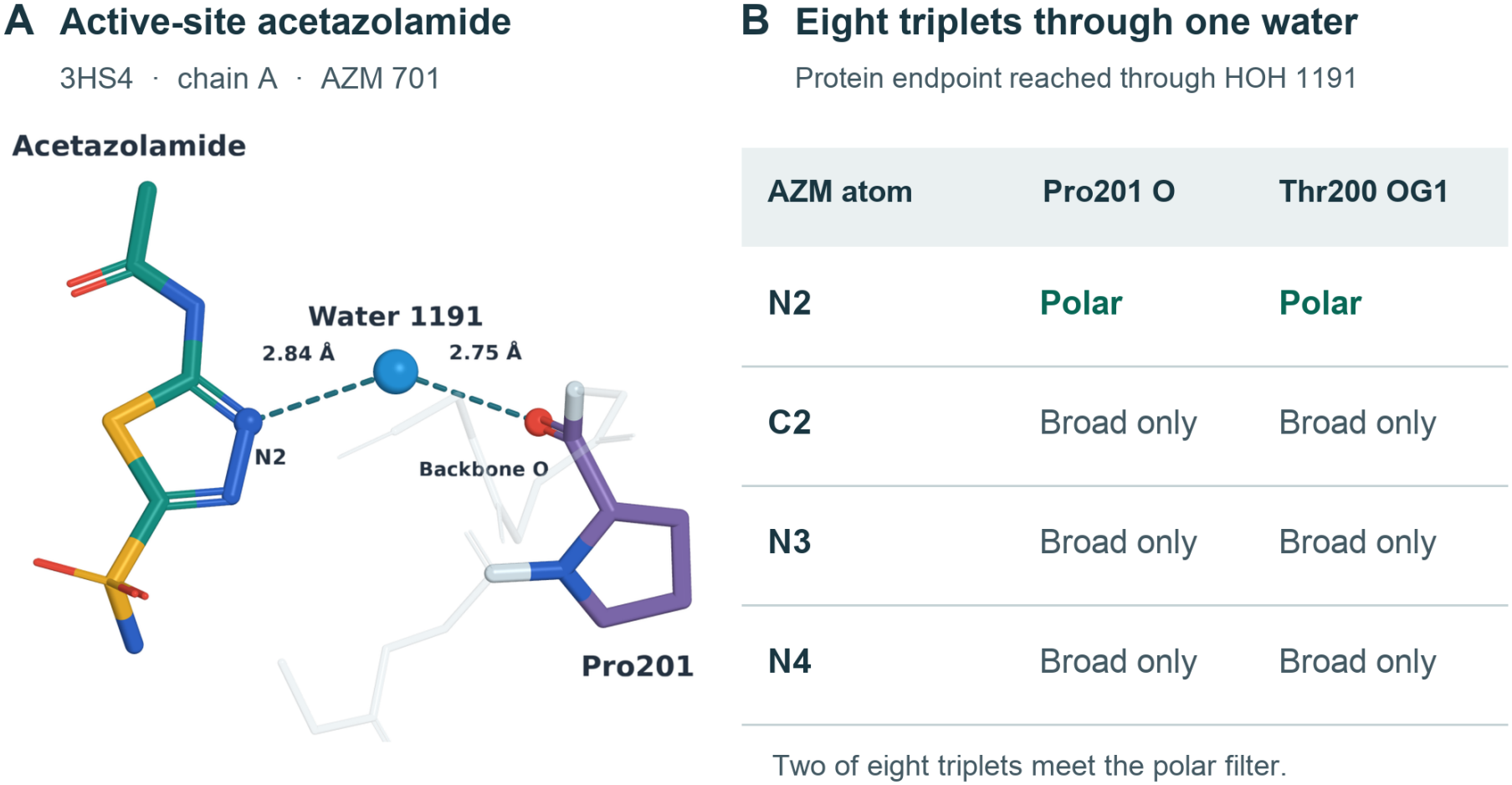
Eight geometric paths through water 1191 in carbonic anhydrase II (3HS4). (A) The native PyMOL view shows acetazolamide N2, water 1191 O and Pro201 O. (B) Each cell represents one exported path. Both N2 paths meet the polar filter and reach different protein atoms (Pro201 O or Thr200 OG1).

Structure Analysis distinguishes aromatic-residue proximity from ring packing. Five aromatic-class residues contact acetazolamide. Using atom and bond definitions from the wwPDB Chemical Component Dictionary [16], StrucMind identifies one complete ligand ring. No ring pair meets the stacking rule. In the estrogen receptor α–estradiol complex 1A52 [17], the ligand and Phe404 ring centers are 5.086 Å apart (interplane angle 74.4°, lateral offset 0.848 Å), consistent with T-shaped packing. Removing a required ring atom withholds the ring assignment while preserving distances among observed atoms.

## Data and software availability

Both StrucMind 0.3.2 plugins are freely available to reviewers and readers from the University of Wyoming ARCC lab website. Download the compiled release for Linux x86-64 or Apple silicon at https://pages.arcc.uwyo.edu/elokely-research/website/assets/downloads/StrucMind_0.3.2_PyMOL_plugins.zip. Both plugins run locally without registration. Installation instructions are in SI S1 and S12. Users may redistribute the unmodified release. The engine source is private.

## Supporting Information

The PDF provides installation steps, contact rules and worked examples. Supporting_Data_and_Sessions.zip contains Data_S1.zip (contact-map tests), Data_S2.zip (ligand environments), Data_S3_DHFR_walkthrough.zip (DHFR comparison), Data_S4_CDK_reconciliation.zip (kinase counts), Linux_validation_0.3.2.zip, Installation_validation_0.3.1.json, Installation_validation_0.3.2.json, General_examples.pse and PyMOL_Sessions_S1_S2.zip.

## Acknowledgment

The authors are responsible for the scientific content of this article, including the analyses, interpretations and conclusions. At the authors’ direction, OpenAI Codex assisted with drafting and revision, plugin implementation and testing.

