## Supplementary material for "StrucMind links protein contacts and ligand environments to atoms in PyMOL": SI document

#### Download StrucMind for PyMOL

Reviewers and readers can download both compiled plugins without signing in. https://pages.arcc.uwyo.edu/elokely-research/website/assets/downloads/StrucMind_0.3.2_PyMOL_plugins.zip

**Examples.** Extract Supporting_Data_and_Sessions.zip from the journal, then unzip the dataset for your example. Walkthrough screenshots are in the StrucMind guide.

**Verify the download.** Compare the command output with the release checksum (SHA-256). Extract the outer ZIP. Keep the plugin ZIPs intact.

**macOS** shasum -a 256 StrucMind_0.3.2_PyMOL_plugins.zip
**Linux** sha256sum StrucMind_0.3.2_PyMOL_plugins.zip

#### Install and verify both plugins

**1. Choose your platform.** For Linux x86-64, follow the tested installation on S12. On Apple silicon use open-source PyMOL 3.1.0 (arm64, Python 3.14, Qt 6) or licensed 3.0.3 under Rosetta (x86_64, Python 3.10, Qt 5). The macOS release is unnotarized and Gatekeeper may block a downloaded copy. Both plugins run locally.

**2. Verify and install on macOS.** From 04_Software, run shasum -a 256 -c SHA256.txt in a terminal. After every file reports OK, keep both ZIPs intact beside Install_StrucMind.py. In PyMOL choose File → Run Script…, open that installer and click Install both plugins. Accept Structure Analysis, then Contact Maps. Save any session, quit and reopen the same PyMOL build.

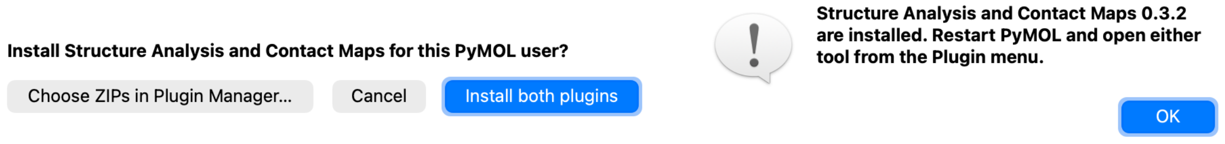


Cropped PyMOL 3.1.0 installer dialogs show the installation choice and completion in a fresh profile. For panel assembly, see the article Acknowledgment.

**3. Check registration.** Open Plugin → Plugin Manager → Installed Plugins. Check for strucmind_papers (Structure Analysis) and strucmind_pymol (Contact Maps), both version 0.3.2.

**4. Test both plugins.** Unzip Data_S3_DHFR_walkthrough.zip. In its folder, run python3 restore_inputs.py (Python 3.8 or later) to retrieve and verify the RCSB coordinates and MTX ligand definition. Follow S10–S11 for the 877 DHFR pairs and 2.392 Å methotrexate contact, or S2 and S4 for your own inputs.

### Compare protein structures on a shared residue axis

Choose Plugin → StrucMind. One input gives one contact map. With multiple inputs, the first input defines a common residue axis, and the Reference and Comparison controls determine the direction of each pairwise contrast.

**Select structures.** Choose Load structures… for PDB or mmCIF coordinates, including compressed files, or Refresh objects for open structures. Select the coordinate state and chain or assembly copies. Choose Add to comparison for each input. Load a deposited biological assembly when the question concerns subunit interfaces.

**Calculate and inspect.** Set the packing cutoff, then choose Calculate contacts. Select Contact type and the neighbor setting. Click a map cell or table row. Focus reference atoms and Focus comparison atoms reveal available atom witnesses. Partner changes… lists the partners gained and lost by each position. Export comparison… writes all statuses for the selected input pair and contact layer.

#### Worked comparison of zinc-containing carbonic anhydrase II

Open sample pair loads chain A of 2CBA as reference and 3HS4 as comparison. With adjacent observed residues included, 1,248 and 1,244 residue pairs meet the 4.5 Å distance cutoff, respectively. After coordinate checks, 1,223 and 1,175 pairs have state status present. Across the union of 1,269 candidates, 19 pairs are gained, 18 lost, 1,134 retained and 98 unresolved (Table S4).

Select the Thr37–Gln255 gain. The Thr37 backbone O to Gln255 NE2 distance is 5.379 Å in 2CBA and 2.697 Å in 3HS4. Each Focus button shows the corresponding atom pair. The gain describes these two deposited coordinates. The same comparison alone cannot assign the structural difference to inhibitor binding.

#### How positions and copies are matched

Identical observed sequences map by position. Other chains use BLOSUM62, gap opening −10, extension −0.5 and free terminal gaps. The method keeps positions shared by inspected optimal alignments. At 33 equally scoring alignments it restricts mapping to consensus positions supported by unique, exact seven-residue anchors.

A compatible copy requires ≥30% identity, ≥50% coverage of the shorter chain and eight aligned positions, or the length of the shorter chain if below eight. Chains shorter than six residues require ≥75% identity. Compatible segment/author-chain identities take precedence. Resolve ambiguous homomers by explicitly selecting copies. Shared insertions require identical internal sequence with adjacent flanking axis positions. Other unmatched sites remain input-specific.

#### Local calculation and versioning

Structure Analysis supplies the local engine for both plugins. The macOS release retains the 0.3.1 scientific engine. Linux uses native x86-64 builds and reads formal-charge information as described on page S12. Map engine_version 0.2.0 identifies the comparison algorithm. Data S1 and S2 retain their run versions. Each platform distributes a readable installer and registration wrappers. Engine source remains private. Both clients verify the compiled runtime against its release manifest. Recipients can inspect the distributed binaries.

### Read the contact rules and coverage calls

For packing, the table includes a pair when an observed heavy-atom distance meets the cutoff in either input. StrucMind then evaluates each state. Present in both gives retained. Absent followed by present gives gained, and the reverse gives lost. A candidate is unresolved whenever either state is unknown. Five carbonic-anhydrase candidates have no final present state despite an observed distance inside the cutoff (Table S4).

An observed atom pair can establish packing in a partially resolved standard residue. A supported absence requires the expected backbone and side-chain heavy atoms in both residues. Alternate conformers, ambiguous mapping and nonstandard chemistry can prevent classification. A missing shared-water observation leaves solvent support unknown. The TSV columns ref_context and target_context give the state-specific explanation.

Table S1. Protein–protein geometric layers in Contact Maps. Distances are inclusive. The packing cutoff is adjustable from 0.1 to 20 Å. The remaining rules are fixed.

| Layer | Positive observation in one coordinate state |
|---|---|
| Packing | Minimum distance between heavy atoms in two protein residues ≤ cutoff (default 4.5 Å). |
| Ionic | Arg NE/NH1/NH2 or Lys NZ to Asp OD1/OD2 or Glu OE1/OE2 ≤4.0 Å. |
| Donor–acceptor | Eligible donor and acceptor heavy atoms ≤3.5 Å. Atom roles come from standard side chains and backbone N (except Pro), O and OXT. Coordinates support a polar contact candidate. |
| Hydrophobic | Nonpolar side-chain carbons in Ala, Val, Leu, Ile, Met, Pro or Phe at ≤4.5 Å. |
| Disulfide geometry | Distinct Cys SG atoms at ≤2.3 Å. |
| Parallel aromatic | Complete Phe, Tyr, His or Trp subrings. Center distance ≤5.5 Å, unsigned plane angle ≤30°. Relative to each plane, height ≥2.5 Å and lateral offset ≤2.5 Å. |
| Cation–π | Lys NZ or Arg CZ (NE, NH1 and NH2 present) to a ring center ≤6.0 Å. Displacement lies within 45° of the normal, with height ≥2.0 Å and offset ≤2.5 Å. |
| Shared water | Polar atoms in two residues are each within 3.5 Å of the same water oxygen. This layer recognizes HOH, WAT and DOD, with oxygen names O, OW or OH2. |

Rows can contribute to multiple layers. In typed layers the closest qualifying protein atom pair is recorded, with ring centers or water identity and leg lengths in the context fields. Disabling neighboring observed residues removes within-chain pairs adjacent in the observed sequence order in either input. Interchain pairs remain eligible.

#### Choose a different pair without changing the shared axis

As a three-input example, Data_S1 includes 4HHB chain A (α), 4HHB chain B (β) and 1HHO chain A (α). The shared axis contains 149 columns. Selecting the two α chains at 4.5 Å yields 20 gains, 17 losses, 627 retained and zero unresolved pairs. Each additional pair with the β chain also includes paralog sequence differences. The three pairwise results share inputs.

### Choose a ligand and measure its pocket

Open Plugin → StrucMind · Structure analysis and select Ligand and pocket. Load structure… accepts PDB or mmCIF coordinates. Refresh objects lists structures already open. Select the object, state, receptor chain(s), ligand instance and alternate. Residue code, author chain, number and segment distinguish repeated ligand copies. Choose Peptide chain for a bound peptide and a separate receptor selection.

The calculation uses occupied heavy-atom coordinates and the common atoms plus one conformer per residue. Automatic choice uses highest mean positive occupancy, then the alternate label to break ties. For a small molecule outside the bundled definitions, choose Load CCD file… and select the matching wwPDB Chemical Component Dictionary (CCD) CIF. Choose Calculate features. Recalculate after changing coordinates, alternate, receptor or ligand selection.

Table S2. Measurements and their biological interpretation.

| View or control | What the estimate describes |
|---|---|
| Measurements and ligand feature atoms | The observed mass center uses natural-abundance elemental weights. The geometric centroid weights atoms equally. Observed mass, radius of gyration and a legacy integer-mass center are separately identified. Show ligand center of mass places a marker in the selected structure frame. |
| Chemical descriptors | For a matched, complete CCD heavy-atom graph, RDKit supplies molecular weight, heavy-atom count, rotatable bonds, topological polar surface area and calculated logP, with reference-graph charge, donor/acceptor counts and atom-resolved features. CCD protonation is the chemical reference. Partial coverage preserves measured coordinate geometry and withholds graph-dependent descriptors. |
| Observed ligand volume | Union of observed heavy-atom van der Waals spheres, using a 0.20 Å ligand-fixed grid and zero probe radius. The result describes this deposited ligand atom set, including an explicitly selected peptide chain. |
| Pocket shells and residue center distances | First shell ≤4.5 Å; second shell >4.5 to ≤8 Å by nearest ligand–residue heavy atoms, with a resolved receptor Cα. Inspect the pair of atoms for each shell member. Composition fractions may overlap. Residue-to-ligand center distances are distinct from the nearest-atom distance. |
| Include native grid cavities | Selected receptor chains define a 1.5 Å grid; candidate connected volumes ≥50 Å³ are reported. The nearest candidate is accepted by ligand-shell overlap with its lining or by centroid distance within 1.2 times the equivalent-sphere radius. Inspect assigned status, lining atoms, enclosure and pocket mouth span. Ligand volume and cavity volume measure different spatial objects. |

Pin as reference stores scalar measurements for this window. Calculate a second case and choose Compare with pinned reference to see comparison minus reference for eligible scalar pairs. Use the recorded selection and coverage to interpret size or shell differences.

As a worked scale check, 65L A301 in CDK5 7VDP has 32 observed heavy atoms, a ligand volume of 335.2 Å³ and 19 first-shell plus 29 additional second-shell residues. Its assigned grid cavity measures 111.4 Å³. Comparing these two volumes as a filling fraction would mix the ligand sphere model with the detector's receptor grid model.

### Trace a ligand atom through water to the receptor

Choose Active-site contacts → Open tutorial structure → Calculate features. The input is model 1 of 3HS4, human carbonic anhydrase II with three deposited acetazolamide (AZM) copies. The tutorial selects active-site AZM A701, segment D, with all 13 heavy atoms. The selected receptor is chain A.

**Direct atom pair.** In Amino acid–ligand atom contacts, find THR199 N1, select Thr199 OG1 and click Show selected evidence. The ligand N1–Thr199 OG1 distance is 2.825 Å. All 94 heavy-atom pairs at ≤4.5 Å are available across 13 residues. The zinc ion at the active site is a distinct nonprotein component and falls outside the amino acid table.

**Shared water.** Reopen the structure-analysis window. Select Ligand–water–amino acid bridges and find AZM701.N2 PRO201.O. The row gives the two distances made by one deposited oxygen. Show selected evidence draws both legs. Ligand N2 lies 2.837 Å from HOH A1191 O, segment G. That oxygen lies 2.754 Å from Pro201 backbone O. It also lies 2.906 Å from Thr200 OG1 in a separate row.

**Count waters and paths separately.** The selected state contains 404 waters. Three contact active-site acetazolamide through 11 ligand–water atom pairs. Only HOH A1191 O also contacts chain A under the bridge rule. Four ligand atoms and two protein atoms give eight triplets through this oxygen. Two pass the polar filter. The 582 protein–water atom pairs cover the full receptor.

#### Water contact rules

HOH, WAT, H2O, DOD, D2O and SOL contribute one observed oxygen per water. Occupancy then deposit serial or atom index resolve alternate sites. A ligand heavy atom within 4.0 Å of the water oxygen and a protein N, O or S within 3.6 Å of the same oxygen define a geometric triplet.

The polar flag requires N or O at both endpoints, both legs ≤3.5 Å and water occupancy ≥0.70. PLIP reports water identities and both distances for its chemically defined water bridges (main text ref. 10). StrucMind exports every geometric triplet with its polar flag, so either set can be counted by distinct water oxygen. Absent water coordinates leave solvent evidence unavailable. An observed solvent set with zero qualifying contacts gives a measured zero. A separate legacy flag marks receptor Cα–water distances ≤6.5 Å. Neither filter measures solvent residence time.

#### Inspect the saved acetazolamide evidence

Unzip Data_S2.zip and open the Data_S2 folder. In 3HS4_AZM701/direct.tsv, find ligand N1 and Thr199 OG1. In bridges.tsv, count the rows, then filter polar_3_5 to True. Group water_index within this selection to count distinct oxygens. Read ligand_water_A and water_protein_A for each retained path. In General_examples.pse, recall Direct_contact and Water_bridge to inspect the atoms.

### Examine a connected water network

Choose Surface-ligand water network → Open tutorial structure → Calculate features. The structure is again 3HS4, now with surface AZM A702, segment E (occupancy 0.70). The Water clusters near the ligand view opens with the ligand scope selected. Select the six-water component and click Show selected evidence. Figure S1 shows how to identify its row.

**A Choose a water component**

3HS4 / acetazolamide A702

| Cluster | Waters | O–O edges | Ligand-near waters |
|---|---|---|---|
| W001 | 6 | 7 | 1 |
| W002 | 2 | 1 | 1 |
| W003 | 1 | 0 | 1 |
| W004 | 1 | 0 | 1 |

Four ligand-connected components

**B Trace its members in the structure**

Six waters · seven O–O edges

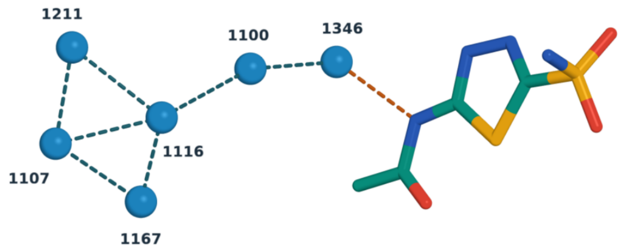


Water 1346 O to acetazolamide N4

**2.514 Å**

Figure S1. Water network adjoining surface acetazolamide A702 (segment E) in 3HS4, model 1. The StrucMind table lists the four components that reach this ligand. The selected component has six waters and seven O–O edges. Only A1346 lies within 4.0 Å of this ligand (N4–O, 2.51 Å). In the PyMOL view, blue spheres mark water oxygens, and teal dashes join pairs within 3.5 Å. For panel assembly, see the article Acknowledgment.

Each node is an observed water with occupancy ≥0.70 within 3.5 Å of a selected receptor N/O atom. Edges join eligible oxygens separated by ≤3.5 Å. The ligand scope retains an entire connected component if at least one member lies within 4.0 Å of any ligand heavy atom. A component can therefore extend beyond the ligand shell.

For this copy, 58 direct ligand–protein atom pairs involve six residues. The four ligand-connected component sizes are 1, 1, 2 and 6. The six-member group contains waters 1100, 1107, 1116, 1167, 1211 and 1346. For the active-site copy A701, the same network rules yield one isolated node, water 1191. Change the scope to All receptor waters to inspect 283 eligible oxygens, 135 edges and 159 components across chain A. These are connected sites in a single deposited coordinate state.

### Separate aromatic proximity from ring geometry

Choose Aromatic packing → Open tutorial structure → Calculate features. This example uses estradiol EST A1, segment C, in the human estrogen receptor α ligand-binding domain (1A52). Ligand aromatic rings lists the atoms defining each ligand ring. Aromatic interaction geometry reports the interacting receptor group, ligand ring, center distance, unsigned plane angle and in-plane offset.

**Inspect the ring pair.** Select Phe404 and choose Show selected evidence. The distance between the ring centers is 5.086 Å, the unsigned interplane angle is 74.4° and the lateral offset is 0.848 Å. The record meets the T-shaped geometry rule. The full ligand contact table contains 112 atom pairs across 17 residues.

**Trace the aromatic cluster.** In Ligand-linked aromatic clusters, select the two-node component, then open Aromatic cluster edges. In this example a single geometric edge joins the ligand and Phe404. With additional geometry-qualified groups, a component can contain protein residues connected indirectly to the ligand.

**Run the comparison case.** Repeat with active-site acetazolamide in 3HS4. Five aromatic-class residues touch the ligand under the heavy-atom cutoff. Its matched CCD provides one ligand ring, with zero qualifying protein–ligand ring pairs. If ligand ring coverage is unavailable, the software withholds the ring count while retaining observed direct distances.

#### Geometry used by Structure Analysis

For a small-molecule ligand, aromatic group membership requires a matched chemical graph and complete observed heavy atoms. Canonical protein Phe, Tyr, Trp and His side chains supply ring templates. A peptide ligand can supply a complete canonical aromatic side chain. A contact to a residue in an aromatic class can involve a backbone or side-chain atom outside that ring.

Stacking candidates require a center distance of 2.0–5.5 Å and an in-plane offset ≤3.0 Å on the recorded reference ring plane. Unsigned angles <30° are parallel; >60° are T-shaped; intermediate angles retain their own category. Cation–π candidates use Arg CZ or Lys NZ within 6.0 Å of a ring center and offset ≤2.0 Å. Small-molecule cations are outside the recognition rule. Each accepted edge records its geometric witness.

The protein-only Contact Maps rule on page S3 uses its own parallel-stacking criteria. Use the module and layer name when quoting a result. The 1A52 record describes geometric packing in one receptor structure; the measured distance and angle do not quantify stabilization energy.

As a separate solvent control, the 1A52 example contains 30 observed waters and zero within 4.0 Å of EST A1. This is an evaluated absence in the specified ligand shell. It differs from a coordinate file with no observed waters, in which ligand hydration cannot be evaluated.

### Export evidence and verify a result

Choose a view, filter the table and select a row. Show selected evidence checks the selected coordinates and draws the recorded atoms or group centers in PyMOL. Reopen the plugin to return to that row. Export displayed table… in Structure Analysis writes the displayed rows to TSV. Save full record… writes the complete analysis to JSON. Contact Maps instead exports the full chosen comparison and layer, including rows hidden by its display filters.

Table S3. Records needed to interpret or repeat a selection.

| Record | How to use it |
|---|---|
| Structure and state | Record PDB accession, coordinate source or assembly, model/state, receptor selection and reference direction. Preserve the input files. The file digest identifies archived bytes; the structure-analysis coordinate digest identifies the submitted atom records in that selection. |
| Atom identity | Retain segment or copy, author chain, residue name and residue identifier (including insertion code), atom name and alternate location. PyMOL atom indices address the currently loaded object; the complete identifiers permit checking a reloaded file. |
| Rule and scope | Report module, layer, contact cutoff, alternate policy and neighbor setting where applicable. State whether water results refer to ligand-connected components or the selected receptor. Report counts as atom pairs, triplets, unique waters, residues, graph edges or components as appropriate. |
| Missing or borderline | Inspect missingness and assignment fields. For packing, rerun a nearby cutoff to assess borderline distances. A zero requires an evaluated selection. An unavailable water count, unassigned cavity or unmatched CCD has a different meaning. |

#### Checks on the released workflows

Independent enumeration recovered the 1,248 and 1,244 geometric candidates in the carbonic-anhydrase maps. Table S4 distinguishes these from state labels after coordinate checks. We remeasured all 94 active-site acetazolamide atom pairs, both Pro201 water-bridge legs and all 13 edges of a receptor water component. Filters and scope were checked in the TSV export.

Both 0.3.2 installation routes passed in the visible menus of open-source PyMOL 3.1.0. After restart, trypsin (2PTN → 3PTB) yielded 1,050 packing assignments and 105 benzamidine A1 atom pairs across 13 residues. Gly219 O–BEN N1 was closest at 2.815 Å. Offscreen PyMOL 3.0.3 File-menu tests covered initial and repeat installation, cancellation and rollback (Installation_validation_0.3.2.json).

In the visible fresh-profile DHFR walkthrough, StrucMind reported 877 packing assignments and 203 methotrexate atom pairs. Independent searches in native PyMOL and Contact Map Explorer reproduced the contacts in each structure (S10, S13). A separate process reopened the saved atom measurement. Earlier tests included mmCIF input, unavailable chemistry, ring geometry, 96 CDK shell witnesses and 35 grid cavities. Halving the volume grid spacing changed ligand volume by less than 0.1%.

### Study examples and archived checks

The Study examples menu revisits the CDK5 networks and opioid atlas that motivated the general tools. These workflows use their own residue axes and units. Their results complement the general exercises on pages S2–S7.

**CDK networks.** Choose 1H4L → 7VDP and Calculate CDK network. Across 277 shared CDK5 positions, the plugin finds 39 Cα contact gains, 27 losses and 592 retained pairs. One gain touches the 4.5 Å shell of inhibitor 65L A301. To inspect the counts originally published using MolScope, choose Published CDK tables… and open the intact supplement by Nael, Alakonda and Elokely (2026, DOI 10.1021/acs.jcim.6c00886.s002). S14 and Data S4 reconcile the counts.

**Opioid atlas.** Choose Opioid atlas cohort → Load / apply filters. The released cohort has 84 independent experimental maps and 3,486 pairwise comparisons. Filter both members by receptor, ligand modality, transducer and state. Compare the independent-map denominator with the number of structure pairs. Look up deposited positions traces an exact generic site to a deposit. Opioid atlas coordinates recomputes ≤8 Å Cα contacts on exact shared TM1–TM7 generic sites.

**Ligand chemistry and peptide controls.** The five core scalar descriptors follow the $\alpha_2$ application. For 6KUX E3F, the 25 observed heavy atoms occupy 279.7 Å$^3$; CCD descriptors give molecular weight 364.51 Da, three rotatable bonds, topological polar surface area 49.85 Å$^2$ and calculated logP 2.43. Peptide-chain examples include the terminal amides of 8F7R and 8F7S; observed volumes are 483.7 and 619.2 Å$^3$.

#### Inspect the earlier archived examples

Data_S1.zip supplies full comparison tables, input digests, mapping fractions, a six-deposit audit and paired workflow grades. PyMOL_Sessions_S1_S2.zip contains replayable μ-opioid receptor and KcsA sessions. From File → Open, select their .pse files and use Scene to recall the supplied states.

Open General_examples.pse and use the Scene menu to replay Direct_contact, Water_bridge, Water_cluster or Aromatic_packing. Data_S2.zip supplies the results for 2CBA, 3HS4 and 1A52, including full_record.json and the nonempty tables. Run python3 restore_inputs.py in the extracted Data_S2 folder to retrieve and verify the RCSB coordinates and the AZM and EST ligand definitions. The run manifest lists evaluated empty layers. Choose Open evidence reader → Open manifest… and open Data_S2/Reader/manifest.json.

Figure S5 traces the μ-opioid receptor comparison to its atoms. For the KcsA assembly (1R3J → 1R3K), Gly79 N in copy C loses Glu71 OE1 in C (4.098 to 5.211 Å) and gains Gly77 O in the same neighboring copy C-3 (5.762 to 4.412 Å). Each row preserves the partner's subunit across both states.

Across six previous deposits, separate coordinate enumeration agreed on all 49,443 protein packing pairs. Rechecking 78,365 layer-specific atom distances and 3,234 both-leg water bridges evaluated coordinate replay. Earlier conventional and StrucMind workflows were equally accurate on 35 sampled μ-receptor sites and 23 KcsA sites (195 partner/copy records). Nael, Alakonda and Elokely describe the kinase study in J. Chem. Inf. Model. (2026, DOI 10.1021/acs.jcim.6c00886) and the in-house StrucMind workflow in the opioid atlas (bioRxiv 2026, DOI 10.64898/2026.08.14.744957).

### Compare two human DHFR structures

**Load and orient the comparison.** Prepare the Data S3 inputs as on S1. In Plugin → StrucMind, choose Load structures… and open Inputs/1DHF.pdb. Select state 1, chain A and Add to comparison. Repeat for Inputs/1U72.pdb. Set Reference to 1DHF and Comparison to 1U72. Select Heavy-atom packing, a cutoff of 4.50 Å and Include neighboring observed residues. Click Calculate contacts to display the map in Figure S2.

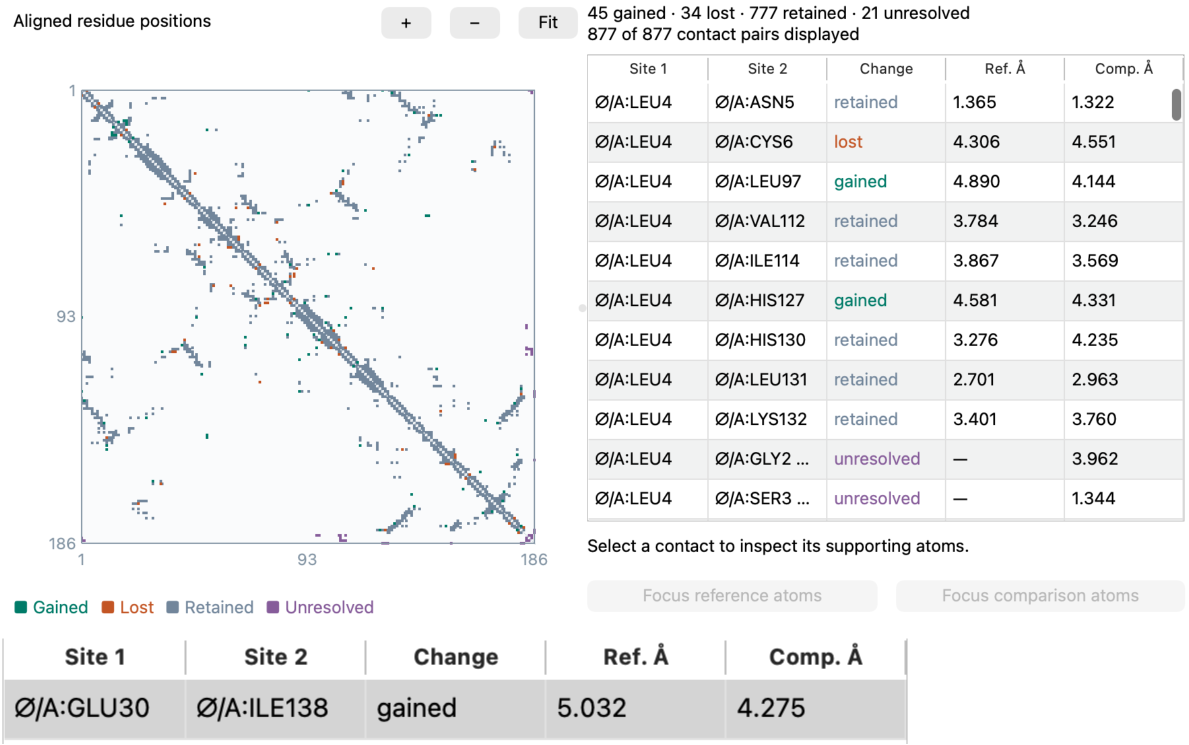


Figure S2. Cropped Contact Maps windows show all 877 packing pairs in the full comparison (top) and the selected gain after filtering (bottom). Each symmetric pair of map cells shares one table row. The 777 retained pairs, 45 gains, 34 losses and 21 unresolved observations include all pairs seen in either map. For panel assembly, see the article Acknowledgment.

**Follow one gain.** Set Show to Gained, search for GLU30 and select Glu30–Ile138. Click Focus reference atoms, then Focus comparison atoms. The closest pair, Glu30 CG–Ile138 CD1, lies 5.032 Å apart in 1DHF and 4.275 Å apart in 1U72. Both reference residues have sufficient heavy-atom coverage to establish absence at 4.5 Å. Export comparison… preserves all 877 rows.

**Inspect an unresolved pair.** Gly2 O–Leu4 N measures 3.962 Å in 1U72. Gly2 lacks coordinates in 1DHF. The export retains the 1U72 distance and records the reference status as unknown. Use the separate map searches and residue matching on S13 to recover all 66 pairs observed only in 1U72.

### Trace methotrexate from a table to its atoms

**Calculate the ligand contacts.** Click Ligand and pocket…, choose the loaded 1U72 object, state 1, receptor Chain A and ligand MTX A188. Retain Auto (occupancy), select Load CCD file… and open Inputs/MTX.cif. Click Calculate features. Choose Amino acid–ligand atom contacts and select the Arg70 NH1–MTX188 O2 row in Figure S3.

View Amino acid–ligand atom contacts | Find a residue, atom, water or cluster

203 of 203 rows. Every observed heavy-atom pair within 4.5 Å is listed. A residue may contribute several different atom pairs.

| Amino acic | Protein aton | Ligand | Ligand aton | Distance Å | Residue clas | Ligand ring aton | Protein ring atom |
|---|---|---|---|---|---|---|---|
| A:ARG70 | NH1 | A:MTX188 | O2 | 2.3923 | polar | No | No |
| A:ARG70 | NH2 | A:MTX188 | O1 | 2.6579 | polar | No | No |

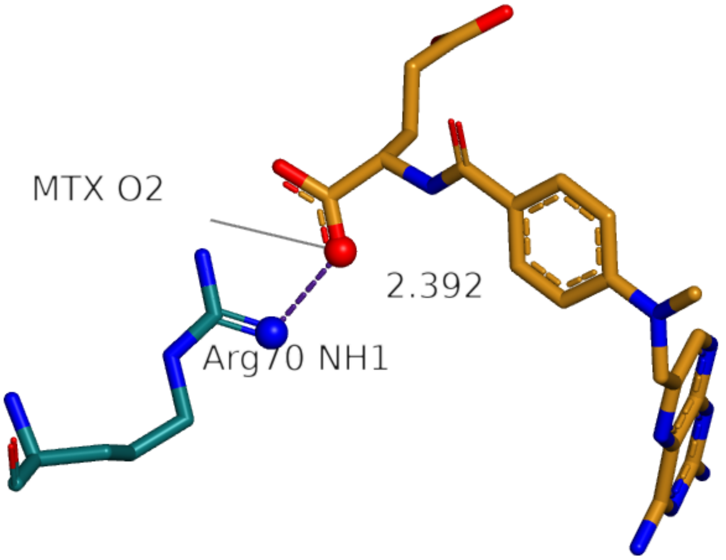


Figure S3. The cropped plugin table shows the selected row among 203 atom pairs (top) and the PyMOL view shows the measured atoms (bottom). Purple dashes join Arg70 NH1 (blue) to methotrexate O2 (red). Arg70 and methotrexate carbons are teal and gold. For panel assembly, see the article Acknowledgment.

**Read and replay the measurement.** Show selected evidence draws the 2.392 Å separation in chain A. This is the shortest observed heavy-atom pair between the selected ligand and protein. The 203 pairs involve 19 distinct amino acids. The distance supports a close polar contact in this model. A binding energy or interaction strength would require other evidence.

**Verify the record.** Use Export displayed table… with the filter cleared, or Save full record… for the complete JSON. In the archived Results folder, the direct-contact TSV retains both atom identities and distance. Open 1U72_MTX_Arg70.pse and recall the MTX_Arg70 scene to revisit the endpoints. Independent enumeration reproduced every contact. A separate PyMOL process reopened the scene and confirmed the 2.392 Å measurement.

### Install and verify the Linux release

Linux was tested on Debian 12 x86-64 with open-source PyMOL 3.1.0, CPython 3.11.18 and Qt 5.15.15. Use X11 or XWayland with OpenGL. Both plugins share a local compiled engine.

#### Install the tested environment

In a terminal opened in 04_Software/Linux_x86_64, run `sha256sum -c SHA256.txt`. When all files report OK, run these commands with micromamba. The conda-forge download requires network access.

```
micromamba create -y -n strucmind032 --file environment-linux-64.txt
micromamba run -n strucmind032 pymol
```

Keep the two ZIPs intact beside Install_StrucMind.py in the Linux folder. In PyMOL, open that installer via File → Run Script…, click Install both plugins and accept both confirmations. Quit and repeat the launch command. Verify both versions as on S1.

#### Reproduce the myoglobin comparison

Unzip Linux_validation_0.3.2.zip. In its folder, run python3 restore_inputs.py to retrieve the RCSB coordinates and HEM ligand definition. In StrucMind → Load structures…, open 1MBD then 1MBC from Inputs. Add state 1, chain A for each. Calculate heavy-atom packing at 4.50 Å with neighbors included. Expect 611 retained, 15 gained, 7 lost and 118 unresolved pairs. Fourteen of the 751 candidates have no final present state (Table S4).

In Plugin → StrucMind · Structure analysis → Ligand and pocket, select 1MBC, state 1, chain A and HEM A155. Load CCD file… → Inputs/HEM.cif, then Calculate features. In Amino acid–ligand atom contacts, expect 235 pairs across 20 residues. Select His93 NE2 and HEM FE, then Show selected evidence (2.192 Å). The HEM dictionary leaves iron charge unknown. Linux preserves the distance and flags charge-dependent chemistry as unavailable.

The install and restart used an unprivileged account and graphical menus on a virtual X11 display in an isolated Debian x86-64 userspace under translation in a Linux virtual machine. The archive includes results, the atom-level scene and verification records.

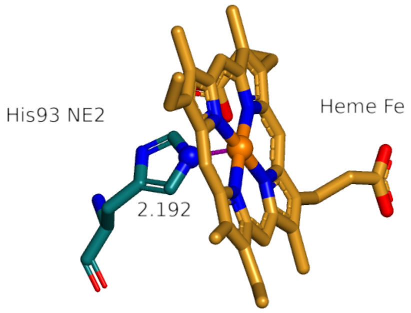


Figure S4. Scripted replay in Linux PyMOL shows His93 NE2 and heme A155 iron in 1MBC. PyMOL recovers the 2.192 Å separation when the saved session is reopened.

### Reproduce the DHFR comparison

Extract Data_S3_DHFR_walkthrough.zip from the Supporting Information and prepare its Inputs folder as on S1. The Comparator folder contains two scripts, tested commands, software versions and expected results. Each script reads the deposited structures directly and can run without installing the StrucMind plugins. See Comparator/README.txt.

Each script enumerated contacts separately in each structure using state 1, chain A, occupied protein heavy atoms, adjacent residues included and a 4.50 Å cutoff. Contact Map Explorer 0.7.0 used ContactFrequency at 0.45 nm. Native PyMOL 3.1.0 used find_pairs in distance mode. Each workflow matched the resulting pairs by deposited chain and residue identifiers. The deposits share a human DHFR sequence and numbering. The 811 reference and 843 comparison pairs yielded 777 shared, 34 reference-only and 66 comparison-only observations. Contact Map Explorer's built-in difference operation restricts comparison to eligible sites in both maps (main text ref. 5).

StrucMind evaluated the 66 comparison-only observations against coordinate coverage. The reference supported absence for 45 pairs. Each of the 21 unresolved pairs involves Val1, Gly2, Ser3 or Asp186, all unobserved in 1DHF. Each row retains the 1U72 atom measurement and the state-specific reason for its assignment. The PyMOL verifier also enumerated 203 protein–methotrexate pairs and reproduced the 2.392 Å Arg70 NH1–MTX O2 minimum.

The coordinate checks use the deposited structures. Data S3 supplies the scripts, settings and expected results. The article Acknowledgment describes assistance with implementation and the coordinate checks.

#### How to read the contact counts

Table S4. Chain A packing at 4.50 Å. Each paired count is given in reference/comparison order. The union counts each residue pair once.

| Structures | Inside cutoff | State present | Union | Neither state present |
|---|---|---|---|---|
| 2CBA → 3HS4 | 1,248 / 1,244 | 1,223 / 1,175 | 1,269 | 5 |
| 1DHF → 1U72 | 811 / 843 | 811 / 843 | 877 | 0 |
| 1MBD → 1MBC | 729 / 737 | 683 / 665 | 751 | 14 |

To reproduce Table S4, open the packing TSV for each comparison in Data S2, Data S3 or the Linux validation ZIP. Count ref_distance and target_distance values at or below 4.50 Å, then separately count present entries in ref_status and target_status. The last column counts rows in which neither status is present. Such rows retain a geometric candidate and its coordinate-quality explanation.

### Reconcile the CDK5 study example

We compared chain A in 1H4L and 7VDP. The 0.3.2 plugin uses the 277 shared canonical CDK5 positions with observed Cα atoms. Among the 37,413 pairs at least four positions apart, a distance ≤8 Å defines contact. A gain touches the inhibitor shell if a heavy atom of either residue lies within 4.5 Å of a heavy atom of 65L A301 in 7VDP. The compiled worker and an independent calculation agree on 592 retained pairs and the changes in Table S5.

Table S5. Change counts under the published analysis and three coordinate calculations. The final column uses each analysis's stated ligand-adjacency rule.

| CDK5 comparison | Gained | Lost | Ligand-adjacent gains |
|---|---|---|---|
| Published Table S8 | 39 | 33 | 13 |
| Plugin, shared canonical positions | 39 | 27 | 1 |
| Observed-sequence alignment | 38 | 33 | 1 |
| Figure-informed reconstruction | 39 | 33 | 1 |

**What changes with the mapping.** A sensitivity calculation aligns the observed chain sequences using BLOSUM62 and affine gap penalties of −10 and −0.5 and measures sequence separation along the aligned rows. It pairs 1H4L Tyr15 with 7VDP Glu12. All six additional losses involve Tyr15 paired with a second residue. Published Figure 3B2 labels the same six partners (Lys33, Arg34, Val35, Arg36, Val44 and Ala48). The aligned-row separation rule also excludes the Leu37–Val44 gain. Assigning Tyr15 to 7VDP Gly11 instead reproduces both published change totals and all ten labeled distance shifts in Figure 3B2 at the precision of the plot. In this figure-informed reconstruction, Tyr15–Phe19 replaces Leu37–Val44 among the 39 gains. Neither Glu12 nor Gly11 represents Tyr15 at the equivalent CDK5 position.

**Which gains touch inhibitor 65L.** Only Ala31–Cys83 meets the explicit inhibitor-shell rule among the 39 canonical gains. Its Cα distance changes from 8.119 to 7.779 Å. The published Figure 3B2 caption identifies Tyr15–Phe19 and Tyr90–Gly96 among its ligand-adjacent gains. 7VDP lists Tyr15 as unobserved. Tyr90–Gly96 gains a Cα contact (10.910 to 7.734 Å). Tyr90 CE1 lies 3.211 Å from O1 of the crystallization additive ethylene glycol (EDO A307). The closest atom pair with inhibitor 65L A301 is Tyr90 N–65L C5 at 6.848 Å, outside the inhibitor shell. The published aggregate and these two examples do not identify the other 11 reported ligand-adjacent pairs.

Extract Data_S4_CDK_reconciliation.zip for the position mapping, changed pairs, alternative shell selections and count audit. Run python3 restore_inputs.py inside CDK_reconciliation to retrieve and verify both structures. Use the canonical row in Table S5 when checking the plugin. Use the published row when citing the earlier study.

### Trace an opioid receptor partner change across cutoffs

We compared chain R of naloxone/Nb6-bound 9PXU with chain R of DAMGO/$G_i$-bound 8EFQ. At 4.5 Å, StrucMind reported 102 gained, 122 lost, 1,170 retained and 58 unresolved packing pairs. Of the unresolved pairs, 31 involved incomplete heavy-atom coverage and 27 an unmapped or unobserved position. Both the ligand and the intracellular partner differ between the complexes.

The Asn88 ND2–Thr120 OG1 distance is 4.823 Å in 9PXU and 3.793 Å in 8EFQ (Figure S5). The pair gains packing contact at each cutoff tested (4.4, 4.5 and 4.6 Å). Both distances exceed the 3.5 Å donor–acceptor cutoff. The assignment therefore depends on which contact layer is selected.

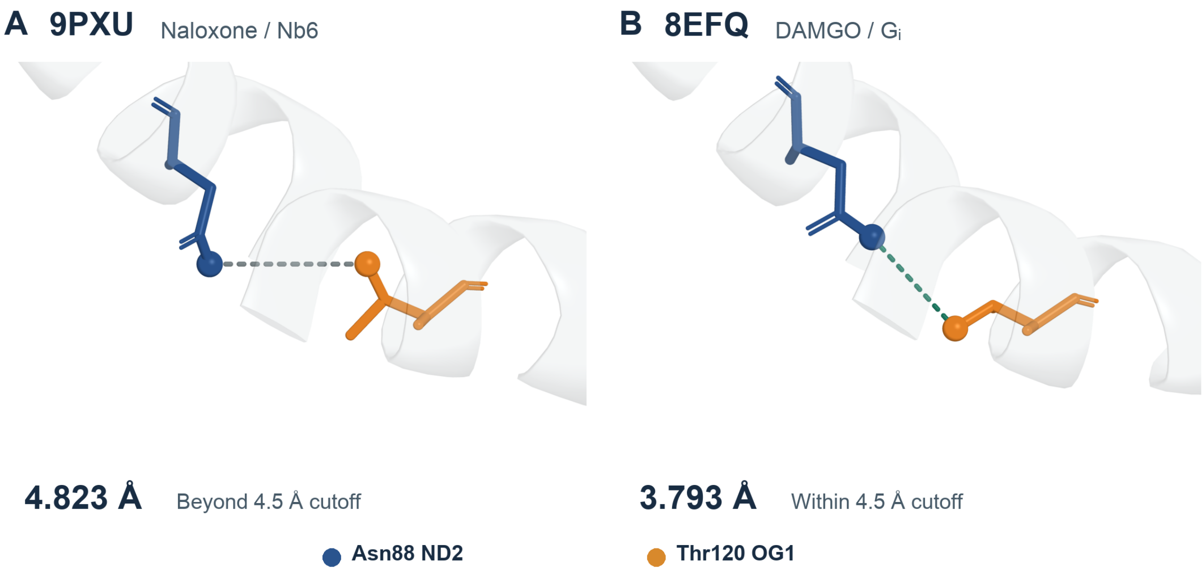


Figure S5. Local Cα alignment orients the two PyMOL views. Each distance was measured in its deposited coordinate frame. The labels identify the same side-chain nitrogen and oxygen in both deposits. For panel assembly, see the article Acknowledgment.

For Asn88–Ser331, the closest heavy-atom distance changes from 2.943 Å (Asn88 ND2–Ser331 O) to 4.575 Å (Asn88 OD1–Ser331 O). The pair is lost at a 4.5 Å cutoff and retained at 4.6 Å. The export identifies the contributing Asn88 atom in each structure.

**Reproduce the comparison.** In Data_S1.zip, inspect OPRM1_Asn88_cutoff_sensitivity.tsv and OPRM1_unresolved_reasons.tsv. Open Session S1 from PyMOL_Sessions_S1_S2.zip and recall the two saved views from the Scene menu.

Structure sources. 9PXU, Khan et al., Nature (2025), DOI 10.1038/s41586-025-09677-6. 8EFQ, Zhuang et al., Cell (2022), DOI 10.1016/j.cell.2022.09.041. The opioid atlas is cited on S9.
